# Crosstalk between invadopodia and the extracellular matrix

**DOI:** 10.1101/2020.02.26.966762

**Authors:** Shinji Iizuka, Ronald P. Leon, Kyle P. Gribbin, Ying Zhang, Jose Navarro, Rebecca Smith, Kaylyn Devlin, Lei G. Wang, Summer L. Gibbs, James Korkola, Xiaolin Nan, Sara A. Courtneidge

## Abstract

The scaffold protein Tks5α is required for invadopodia-mediated cancer invasion both *in vitro* and *in vivo*. We have previously also revealed a role for Tks5 in tumor cell growth using three-dimensional (3D) culture model systems and mouse transplantation experiments. Here we use both 3D and high-density fibrillar collagen (HDFC) culture to demonstrate that native type I collagen, but not a form lacking the telopeptides, stimulated Tks5-dependent growth, which was dependent on the DDR collagen receptors. We used microenvironmental microarray (MEMA) technology to determine that laminin, collagen I, fibronectin and tropoelastin also stimulated invadopodia formation. A Tks5α-specific monoclonal antibody revealed its expression both on microtubules and at invadopodia. High- and super-resolution microscopy of cells in and on collagen was then used to place Tks5α at the base of invadopodia, separated from much of the actin and cortactin, but coincident with both matrix metalloprotease and cathepsin proteolytic activity. Inhibition of the Src family kinases, cathepsins or metalloproteases all reduced invadopodia length but each had distinct effects on Tks5α localization. These studies highlight the crosstalk between invadopodia and extracellular matrix components, and reveal the invadopodium to be a spatially complex structure.

## INTRODUCTION

Cancer invasion, the ability of cancer cells to infiltrate into and through extracellular matrix (ECM), has long been associated with tumor metastasis (Hanahan and Weinberg, 2011). Increasing evidence suggests that the same mechanisms that allow invasive capacity also promote tumor growth during cancer progression. This may be because the ECM acts as a physical barrier to tumor growth, which cancer cells destroy by activating pericellular proteases (Lu et al., 2012), Alternatively, these same pericellular proteases can release latent growth factors and cytokines in the matrix that promote tumor growth and angiogenesis (Chang and Werb, 2001). In either case, tumor growth and tumor metastasis are interlinked.

In many types of cancer, degradation of ECM has been linked to the formation of invadopodia (Murphy and Courtneidge, 2011; Weaver, 2008b), actin-rich plasma membrane protrusions coordinate the cytoskeleton with pericellular proteolytic activity. Since their discovery, invadopodia have been considered characteristic structures of invasive disease. Aside from an array of actin-regulatory proteins and signaling molecules, invadopodia also contain several classes of protease, including members of the metalloprotease superfamily (Chang and Werb, 2001; Seals and Courtneidge, 2003), serine proteases such as fibroblast activation protein (FAP) (Monsky et al., 1994) and urokinase-type plasminogen activator (uPA) and its receptor (uPAR) (Artym et al., 2002), and cysteine proteases such as cathepsins (Tu et al., 2008). Despite our steadily increasing knowledge of the composition and functions of invadopodia *in vitro*, much less is known about their *in vivo* presence and functional relevance *in vivo*. However, related podosome structures have been observed in motile vascular smooth muscle cells and neural crest stem cells *in vivo* (Murphy et al., 2011; Quintavalle et al., 2010), and genetically-defined invadopodia have been observed during intra- and extra-vasation in animal models (Gligorijevic et al., 2014; Leong et al., 2014) as well as in freshly explanted primary human tumors (Weaver, 2008a).

Almost all of the components of invadopodia are also found in other actin-based structures such as adhesion complexes and protrusions (Murphy and Courtneidge, 2011). However, the Tks5 adaptor protein (SH3PXD2A) has a more restricted subcellular distribution, and is found predominantly at invadopodia (Saini and Courtneidge, 2018). Furthermore, knockdown of Tks5 prevents invadopodia formation and function (Abram et al., 2003; Buschman et al., 2009; Iizuka et al., 2016; Seals et al., 2005). Conversely, enforced expression of the longest, α isoform of Tks5 in non-invasive cancer cells lacking native expression results in the formation of invadopodia (Seals et al., 2005). Together these data support the conclusion that Tks5α is an obligate invadopodia scaffold, and its expression can be used as a surrogate to study invadopodia-based invasive capacity.

The Tks5 and related Tks4 adaptors are large scaffolding proteins composed of phosphatidylinositol lipid binding PX domains followed by 5 or 4 SH3 domains respectively. Both adaptors are Src substrates, and also contain multiple serine/threonine phosphorylation motifs. While we do not yet have a full understanding of all binding partners of the Tks adaptors, they have been implicated in reactive oxygen species (ROS) production (Diaz et al., 2009) and have been shown to interact with actin-remodeling proteins (Crimaldi et al., 2009; Oikawa et al., 2008; Stylli et al., 2009), pericellular proteases (Buschman et al., 2009), as well as the small GTPase Rab40b (Jacob et al., 2016), the guanine exchange factor Sos1, and the membrane remodeling GTPase dynamin (Rufer et al., 2009). The Oikawa lab has shown that invadopodium formation is initiated near focal adhesions by the binding of the Tks5α PX domain to PI3,4P_2_ (Oikawa et al., 2008). This model was refined by Condeelis *et al.*, who showed that small actin foci form in the absence of Tks5 (likely following Src phosphorylation of cortactin), and that Tks5 acts to stabilize these nascent protrusions (Sharma et al., 2013). However, few studies to date have examined the structure and function of invadopodia formed *in vitro* in response to extracellular matrix, where their full functionality is observed. Here we describe such experiments.

## MATERIALS AND METHODS

### Cell lines

The luciferase-expressing human breast cancer cell line MDA-MB-231-Luc was obtained from Xenogen. The Src-transformed NIH-3T3 (Src3T3) cells have been described previously (Seals et al., 2005). The cell lines were routinely cultured in Dulbecco’s minimal essential medium (DMEM) supplemented with 10% Fetal Bovine Serum (FBS) and a cocktail of penicillin and streptomycin (P/S) (Gibco). The human breast cancer cell lines, Hs578t and HCC1806, UACC812, SUM44PE, SUM52PE, HCC1187, SUM1315MO2, HCC1143, HCC1937, BT474 and HCC3153 were kindly provided by Dr. Joe Gray (Oregon Health and Science University [OHSU], USA). The culture medium used: SUM52PE (HAMS F12, 5% FBS, 1M HEPES, 1mg/mL hydrocortisone, 10mg/mL Insulin), HCC1187 (RPMI, 10% FBS), SUM1315MO2 (HAMS F12, 10ug/mL EGF, 5% FBS, 10mM HEPES, 5ug/mL insulin), HCC1143 (RPMI, 10% FBS), HCC1937 (RPMI, 10% FBS), BT474 (DMEM, 10%FBS) and HCC3153 (DMEM, 10%FBS).

### DNA constructs and shRNAs

Human Tks5α was cloned into pCDH-CMV-MCS-EF1-Puro (Addgene) with a GFP sequence fused at the C-terminus. The shRNAs pLKO.1 lentiviral plasmids used for scrambled and human Tks5 knockdown were described previously (Blouw et al., 2015). The clones used were TRCN136336 (D5), TRCN0000136014 (D6) and TRCN0000136512 (D7). The shRNAs pLKO.1 lentiviral plasmids used for DDR1 and DDR2 knockdown were: TRC000121082 and TRCN0000361395. The plasmids encoding cortactin (#26722), N-WASP (#54199) and fascin (#54094) were obtained from Addgene. Each insert was cloned into the lentiviral plasmid pCDH-CMV-MCS-EF1-Puro with GFP fusion protein at C-terminus.

### Tks5 antibodies design and production

The antibodies for Tks5 (Rabbit monoclonal, F4 and mouse monoclonal, G6) were generated by Abcam/Epitomics (F4) or by the Oregon Stem Cell Center, Monoclonal Antibody Core Facility at OHSU (G6), and validated by the Courtneidge laboratory. The following primer sets were used for cloning into pGEX-4T1 to produce the immunogens:

PX domain (F4)-Forward: CGGGATCCATGCTCGCCTACTGCGTGCAG

PX domain (F4)-Reverse: CCGCTCGAGCTACTCTTTTGGAGGGTTGACATC

SH-linker (G6)-Forward: CTTCAGAGGATGTGGCCCTG

SH-linker (G6)-Reverse: CCTCACTCTTGGAGCCCTTG

For Protein production: Appropriate DNA constructs were transformed into BL121 E.coli and a single colony was selected and grown overnight at 37 °C in 5 mL of LB medium with ampicillin. The overnight culture was then added to 1 L of LB-ampicillin pre-warmed to 25 °C, grown in a shaking incubator at 25 °C until OD600 was reached 0.5. Follow this, IPTG was added to a final concentration of 0.1mM and cultured overnight at 25 °C. The cells were pelleted in a centrifuge and lysed with lysis buffer (20mM Tris pH 7.5, 1% Triton, 1mg/mL Lysozyme, 5mM DTT and protease inhibitor cocktail, Roche #11-836-170 001) for 1 hour at 4 °C. Lysed cells were placed in pre-cooled tubes in an ice bath, and sonicated at 50% power with Sonic Dismembrator (Model 705, Fisher Scientific) with a micro-tip for 30 seconds, in 5 cycles. Lysates were centrifuged at 10,000xg for 5 minutes at 4 °C and the pellet was discarded. 5 mL of glutathione sepharose 4B was prepared according to manufacturer’s instructions (GE #17-0756-01) and added to the lysates, which were incubated at 4 °C on a rocking platform for 3 hours. The lysates were centrifuged at 500xg for 5 min at 4 °C and was washed in 50 mL of ice-cold 1X PBS for a total of three washes then re-suspended in a final volume of 20 mL of ice-cold PBS. An appropriate size of chromatography column (R&D) was loaded with chilled sepharose beads at 4°C and allowed to settle. Lysate solution was added to the column and allowed to pass through the column by gravity. 5 mL of elution buffer (50mM Tris HCL, 25mM glutathione, pH 8.0) was added to the beads, and this process was repeated for a total of 12-18 cycles. Eluates were collected at each cycle and kept on ice. To determine which elution contained the protein, samples of each elution were loaded onto a poly-acrylamide gel which was stained by coomassie-blue followed by destain (Thermo Fisher Scientific). Eluates that contained the appropriate proteins were pooled together in a pre-chilled tube on ice. Pooled eluates were concentrated in an Amicon Ultra (Millipore) centrifuge unit. Purified lysates were immediately frozen at −80 °C to maintain integrity of proteins. GST tagged proteins were verified by immunoblot with an anti-GST antibody (#71-7500, Zymed). Purified lysates were then sent to Abcam/Epitomics (F4), or to the Oregon Stem Cell Center, Monoclonal Antibody Core Facility at OHSU (G6) for production of monoclonal antibodies.

Immunizations for monoclonal antibody production at OHSU was conducted under a protocol approved by the OHSU Institutional Animal Care and Use Committee. Balb/C mice received multiple IP immunizations with the Tks5 protein. Four days after the final boost, mice were humanly euthanized using CO2 and their spleens were harvested. Splenocytes were fused with SP2/0 Ag14 myeloma cells (Kohler and Milstein, 1975) and hybrid cells were selected by growth in methylcellulose-containing HAT medium (Stem Cell Technologies). Approximately 600 clones were isolated and transferred to liquid HT medium in 96-well plates. Supernatants from these wells were initially screened for Tks5-specific monoclonal antibodies by ELISA.

### Antibodies and reagents for immunoblotting and staining

Antibodies used for immunoblotting were: F4 (1:10), G6 (1:500) and tubulin (T6557, Sigma, 1:3000). Antibodies used for immunofluorescence staining were: F4 (1:10), G6 (1:250) and tubulin (6074, 1:250, Sigma). Fluorescently labeled phalloidin (Alexa Fluor-350, −488, −568 or −647, Thermo Fisher Scientific) was used for actin staining. Hoechst (1:4000, Thermo Fisher Scientific) was used for nuclear staining.

### Invadopodia assay on high-dense fibrillar type I collagen (HDFC)

Invadopodia staining was performed as previously described (Iizuka et al., 2016). Briefly, cells were grown on glass coverslips with or without collagen and fixed with 4% paraformaldehyde/PBS (Electron Microscopy Sciences). For the invadopodia assay on collagen-coated coverslips, HDFC was prepared according to the original protocol reported (Artym, 2016; Artym et al., 2015). Briefly, 18 mm coverslips were pre-chilled on ice and coated with 10 µl ice-cold neutralized collagen (#35429, Corning). The pipette tip was used to spread the collagen evenly on the glass surface and the coated coverslips were left on ice for 10 min to facilitate flattening of the collagen. The layer of collagen was polymerized into a fibrillar meshwork at 37 °C for 30 min, followed by centrifugation at 3,500 g for 20 min. After fixation and permeabilization with 0.1% Triton X-100/PBS for 15 min, the cells were blocked by 5% BSA in PBS with 5% goat-serum for 1 hour at room temperature (RT) and incubated with primary antibodies for 90 min at RT (or overnight at 4 °C). The cells were washed and incubated with Alexa Fluor-conjugated secondary antibodies and phalloidin. Confocal images were collected using a laser-scanning confocal microscope LSM880 equipped with AiryScan (Carl Zeiss). Images were transferred to Imaris™ (Bitplane) which is a multidimensional analysis program based on the fluorescence intensity data to create 3D view of invadopodia and quantify invadopodia length, which was measured by Imaris software using the polygon scaling tool in Slice view.

### 2D/3D proliferation assays

Type I collagen 3D cultures were performed as described previously (Iizuka et al., 2016). For collagen 3D proliferation assay, briefly rat-tail type I collagen (#08-115, Sigma, Millipore) was prepared to a final concentration of 2.1 mg/ml, and 2,500 cells were added to the collagen mix before gelling. For atelocollagen proliferation assay, rat-tail type I collagen or atelocollagen (##602, Yo Proteins) was prepared to a final concentration of 2.1 mg/ml and added to multi-well plate for 1hr in a CO_2_ incubator. Spread cells were grown in DMEM containing 10% FBS. The matrix was dissolved with 2 mg/ml collagenase type2 (#LS004176, Worthington Biochemical corporation) and cell numbers were quantified using hemocytometry. HDFC was prepared according to original protocol reported in (Artym, 2016; Artym et al., 2015) and described briefly above. The HDFC was dissolved with a 1:1 mixture of 2 mg/ml collagenase type2 and trypsin (Gibco), and cell numbers were quantified using hemocytometry. For the growth assay in Matrigel (BD Biosciences), briefly, the cells were added to Matrigel (1:1 dilution with serum-free medium) on ice before gelling in a CO_2_ incubator for 30 min. The cells in Matrigel were cultured in DMEM containing 10% FBS. The matrix was dissolved with cold PBS and cell numbers were quantified by hemocytometry.

### Invasion and growth assay and invadopodia staining in spheroids

3D spheroid cultures were performed as described previously (Kelm et al., 2003). Briefly, spheroids of MDA-MB-231 cells were prepared in hanging droplets with 2,000 cells in 10 µl of 20% FBS containing DMEM for 3 days. The spheroids were embedded in 2.1 mg/ml of rat type I collagen (#354263, Corning) and incubated with DMEM containing 10%FBS for 2 days. Spheroids were fixed with 4% PFA and washed with PBS for three times. After permeabilization with 0.1% Triton X-100/PBS for 15 min, the samples were stained by Alexa Fluor-568-phalloidin to visualized the entire spheroid in 3D collagen. Imaging was performed on Zeiss/Yokogawa CSU-X1 spinning disk confocal microscope and the z-section with the maximum size of actin stained spheroids were collected and measured for the “spheroid size (actin intensity)”. For invadopodia visualization, spheroids were stained by Alexa Fluor-568-phalloidin and Hoechst (1:4000, Thermo Fisher Scientific, used for nuclear staining). Imaging was performed on a laser-scanning confocal microscope LSM880 equipped with AiryScan. Inhibitors used for the spheroid growth assay were: GM6001 (EMD Millipore), CA074 (Vergent Biosciences) and DDR1-1N-1 (Selleck Chemicals).

### Microenvironmental Microarray (MEMA)

MEMAs were produced as described previously (Smith et al., 2019; Watson et al., 2018) with several minor modifications. The same set of 45 ECM or ECM combinations that we previously reported were plated into the bottom of 96 well plates (Ibidi). The plates were left desiccated for several days at room temperature, then 3000 MDA-MB-231 cells containing Tks5αGFP were plated in 100 µl DMEM containing 10% FBS. After overnight adhesion in a 37 °C incubator with 5% CO_2_, an additional 100 µl of DMEM containing PBS control, epidermal growth factor (EGF), fibroblast growth factor 2 (FGF2), or hepatocyte growth factor (HGF) was added to give final concentrations of 10 ng/ml EGF and FGF2 and 40 ng/ml of HGF. The cells were left to grow for 24h, then fixed with 4% PFA and washed with PBS three times. The cells on MEMA were stained with Alexa Fluor-568-phalloidin to visualize the invadopodia, which were also marked with Tks5αGFP, where double positive delineated invadopodia). Hoechst was used for staining of cell nuclei. Imaging was performed on laser-scanning confocal microscope LSM880 equipped with AiryScan.

### Super resolution microscopy

MDA-MB-231 cells with Tks5αGFP were cultured on HDFC prepared on #1.5 coverglass for 2 days before stained for F-actin (phalloidin-Alexa Fluor-647) and Tks5αGFP (GFP-nanobody, ChromoTek GFP Vhh, # gt-250) using manufacturer recommended procedures. The GFP-nanobody was conjugated to an oligonucleotide (‘P1’ docking strand) using published procedures (Schnitzbauer et al., 2017). After staining, the cells were imaged sequentially, first with DNA-PAINT (using a CF660R conjugated ‘P1’ imager strand) in buffer C, which contains 1x PBS supplemented with 500 mM NaCl. Each time after a cell was imaged with DNA-PAINT, the buffer was replaced with 1x PBS supplemented with an oxygen scavenging mixture (comprising glucose oxidase, catalase, and glucose) and 50 mM 2-mercaptoethylamine (MEA) for STORM imaging (van de Linde et al., 2011). For both DNA-PAINT and STORM imaging, a weak cylindrical lens (f=1000 mm) was inserted in front of the detector (Andor iXon Ultra 897) for 3D single-molecule localization (Huang et al., 2008). DNA-PAINT and STORM imaging were performed on a custom setup as described previously (Creech et al., 2017), and 3D localizations were done using ThunderSTORM (Ovesny et al., 2014).

### Activity-base probes (MMP-ABP, CTS-ABP)

Cathepsin activity-base probe (CTS-ABP) were purchased from Vergent Bioscience and imaging was performed according to manufacturer’s instructions.

Chemical synthesis of MMP activity-based probe (MMP-ABP): SiTMR (Lukinavicius et al., 2013), 3-azido-1-propanamine (Hannant et al., 2010), and HxBP (Saghatelian et al., 2004) were synthesized following published protocols. To a solution of SiTMR (200 mg, 423 µmol) in DCM (10 mL) at room temperature, DiPEA (442 μL, 2.54 mmol, 6 eq) was added. The reaction was stirred for 10 mins before HOBt (71 mg, 508 μmol, 1.2 eq), 3-azido-1-propanamine (85 mg, 846 μmol, 2 eq) and EDC (97 mg, 508 μmol, 1.2 eq) were added sequentially. The reaction mixture was covered from light and stirred overnight. The reaction was subsequently added saturated NaHCO_3_ and extracted with DCM (3 × 50 mL). The combined organic layers were rinsed with brine, dried over anhydrous Na_2_SO_4_, and the solvent was removed using a rotary evaporator. The residue was purified by flash column chromatography with silica gel, using DCM/MeOH as eluent to give compound SiTMR-N_3_ (173 mg, 74%) as a pale green solid. Under N_2_, compound HxBP (4.2 mg, 7.84 μmol) and SiTMR-N_3_ (5 mg, 9.02 μmol) were suspended in 2 ml degassed DMSO/H_2_O (1/1, v/v). Freshly prepared 100 mM sodium ascorbate solution (95 μL) and copper sulfate (CuSO_4_) solution (40 μL) were added to the solution. The resulting reaction mixture was stirred at room temperature overnight and purified using preparative high-performance liquid chromatography (HPLC, Agilent 1250 Infinity HPLC) with a C18 column (150 × 21.2 mm). The sample was eluted using solvents A: water-formic acid (99.9:0.1, v/v) and B: acetonitrile-formic acid (99.9:0.1, v/v), with the gradient increased from 10% B to 90% B over 60 mins at a flowrate of 10 mL/min. The fractions containing product were frozen and lyophilized to afford MMP-ABP (4.1 mg, 48%) as a blue solid. HRMS (ESI-TOF) m/z [M + 2H]^2+^ calculated for C_60_H_71_N_9_O_9_Si_2_ 545.7645, found 545.7658. Synthetic route, structure and purity confirmation of MMP-ABP is shown in **Supplementary Figure S1**.

## RESULTS AND DISCUSSION

### New reagents to study Tks5 isoforms

There are multiple isoforms of Tks5, generated by alternative promoter usage (Cejudo-Martin et al., 2014; Li et al., 2013). The full length (α) isoform contains the amino terminal PX domain (Figure 1A) required for phosphatidylinositol lipid binding and the formation of invadopodia (Seals et al., 2005), while the shorter isoforms (β and short) lack this domain. The Tks5-specific antibodies we generated previously (Lock et al., 1998) were raised against the first and third SH3 domains of the mouse protein. As a result, this antibody could not distinguish between the α and β isoforms nor reliably between Tks4 and Tks5; they also did provide high fidelity staining in human cells. To overcome these limitations, we generated new Tks5 antibodies to assist in the study of invadopodia. Of these, F4 is a rabbit monoclonal antibody (produced in collaboration with Epitomics, Inc.) raised against an epitope in the PX domain of Tks5α, and G6 is a mouse monoclonal antibody (produced in collaboration with the Flow Cytometry Shared Resource at OHSU) raised against an epitope in one of the unique linker domains of Tks5. The specificity of these antibodies in immunoblotting is demonstrated in Figure 1B. G6 recognizes both full-length and PX domain-deleted forms of Tks5 (e.g., the α and β isoforms) in human cells, whereas F4 recognizes only the full-length α isoform of Tks5. Neither antibody recognized the related Tks4 protein.

**Figure 1.**
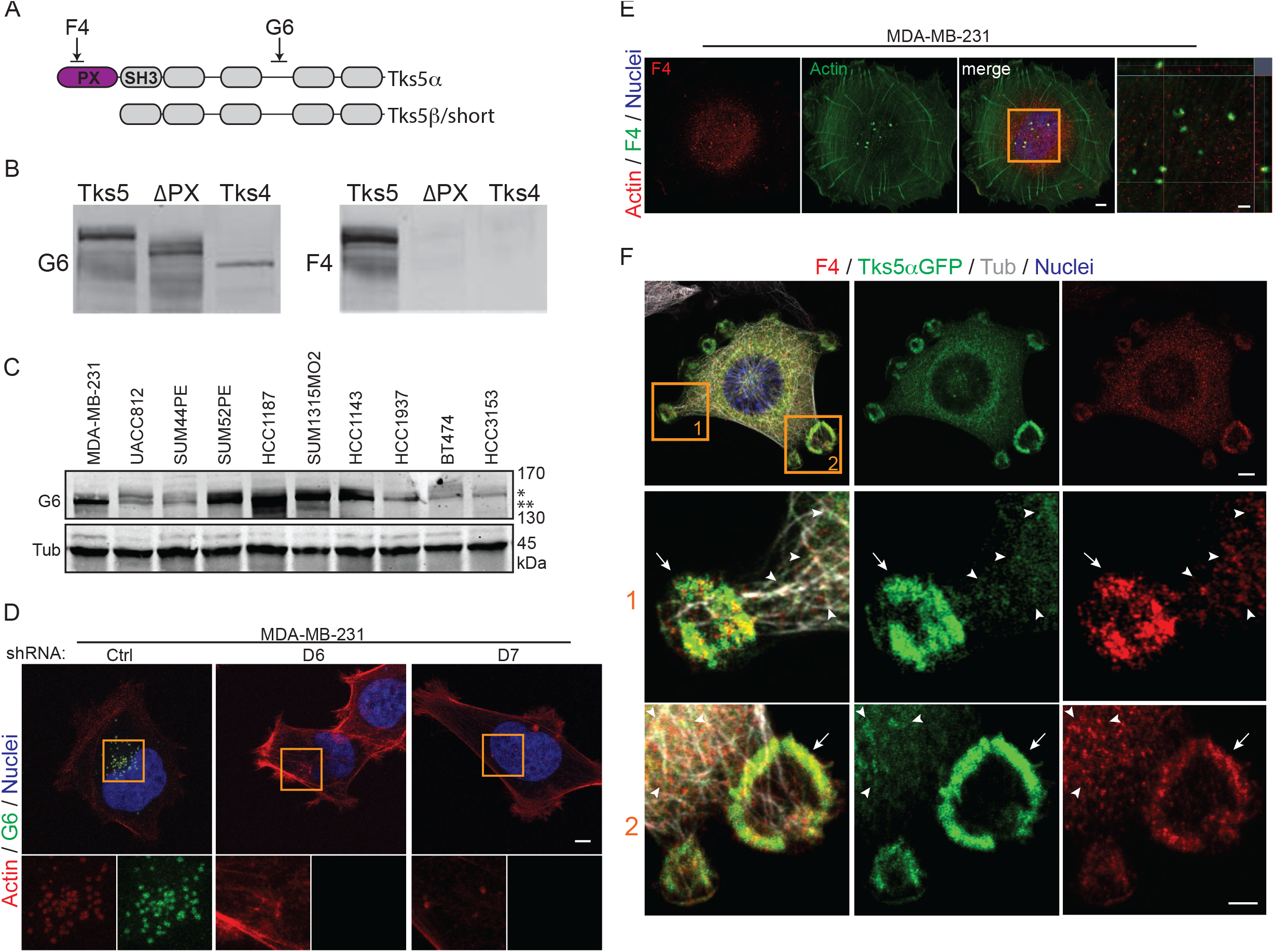
Tks5 antibodies against PX domain (F4) and linker region (G6). **A** Schematic view of Tks5 isoforms and antibody epitope regions. **B** Antibody specificity analyzed by immunoblotting on extracts from 293T cells transiently expressing full-length of Tks5α (Tks5), Tks5α with PX domain deletion (! PX) or full-length of Tks4 (Tks4). **C** Tks5 protein levels analyzed by immunoblotting on human breast cancer cell lines. Asterisks show predicted size of Tks5α (*) and Tks5β/short (**). Tubulin is shown as a loading control. **D and E** Validation of antibodies by immunofluorescence. (D) All Tks5 isoforms were stained by G6 in MDA-MB-231 cells with shRNA-scrambled (Ctrl), shRNA-Tks5 (D6) or shRNA-Tks5 (D7). Invadopodia were visualized by Tks5 (G6, green) and actin (phalloidin, red). (E) Tks5α was stained by F4 in Src-transformed NIH3T3 cells with Tks5αGFP overexpression. Magnified areas (1 and 2, orange square in top) were shown. Arrows show Tks5 accumulation at invadopodia. Arrowheads show Tks5α signals on microtubules in cytoplasmic region. **F** Invadopodia were stained for Tks5 (G6 or F4, red) and actin (phalloidin, green) in MDA-MB-231 cells. Magnified orthogonal views (orange square) were shown in left. Hoechst to denote nuclei of cells in the images. Scale bars, 5 μm (D, E left, F top) and 2 μm (E right, F bottom).

Given our interest in the role of invadopodia in breast cancer, we next used the G6 (“panTks5”) antibody to profile a series of invasive human breast cancer cell lines. All tested cell lines expressed predominantly Tks5α, with low levels of Tks5β and short also detected in some cell lines (Figure 1C). We examined expression by immunofluorescence in one of these cell lines, MDA-MB-231. Focusing on the ventral surface of the cell, we noted co-localization of G6 with the puncta of F-actin characteristic of invadopodia, as expected, whereas no staining was seen if Tks5 expression was reduced by RNA interference (Figure 1D). The F4 antibody also showed specific staining at invadopodia in these cells (Figure 1E). Thus, we conclude that these antibodies are suitable for both immunoblotting and immunocytochemistry. We also found them to provide suitable utility for immunoprecipitation and immunohistochemistry (data not shown). We next used F4 for high-resolution imaging of Tks5α in Src-transformed mouse fibroblasts (Src-3T3), a workhorse for studies on invasive behavior in which the invadopodia form into characteristic rosettes (Chen, 1989; Tarone et al., 1985). Here too we observed Tks5α co-localization with F-actin in the rosettes (Figure 1F). However, in these cells, the relative lack of actin stress fibers, coupled with our use of high-resolution microscopy, also allowed us to observe punctate staining of Tks5α in the cytoplasm. Interestingly, in the cytoplasm Tks5α staining was coincident with tubulin, but not actin, suggesting that Tks5α is localized to microtubules as well as invadopodia. Similar distribution was observed when Tks5α fused with GFP or mCherry was expressed in the cells. This analysis also revealed that the microtubules terminated near the rosettes, in keeping with other reports on podosomes and invadopodia (Linder et al., 2011; Luxenburg et al., 2007; Schoumacher et al., 2010). These data suggested the trafficking of Tks5α on microtubules, which indeed we have now observed (studies to be reported elsewhere).

### Morphology and growth of breast cancer cells with and without Tks5

The ECM, particularly collagen-I, can control cell survival, proliferation, migration and invasion. In cancer, collagen-I had been thought to be growth suppressive (Henriet et al., 2000), but this view is now more nuanced (Keely, 2011; Pickup et al., 2014). For example, increased collagen-I density and changed ECM architecture promote tumor proliferation and metastasis and are linked to a worse clinical outcome (Schedin and Keely, 2011). In addition, ECM rigidity has been shown to promote invadopodia formation (Parekh et al., 2011) and high-density fibrillar collagen (HDFC) in particular increased invadopodia formation in the MDA-MB-231 breast cancer cell line (Artym et al., 2015).

We have previously published that Tks5 knockdown is not required for growth of either breast cancer or melanoma cells on tissue culture plastic, but has an inhibitory effect when cells are placed in a cross-linked matrix of type I collagen (collagen-I) (Blouw et al., 2015; Iizuka et al., 2016). An example of this finding is shown in Figure 2A. Over the course of this five-day assay, control cells increased in number approximately 10-fold, but knockdown of Tks5 with two different shRNAs had a marked inhibitory effect. Interestingly, the basement membrane surrogate Matrigel stimulated growth only three-fold over the same time period, although Tks5 knockdown was still inhibitory. We next wanted to explore whether the ability of collagen-I to form cross-links was important for the observed growth stimulation. To do this we compared native collagen to a pepsinized version which lacked the telopeptides where cross-linking occurred (Figure 2B). We found that, compared to controls, atelo-collagen only poorly stimulated growth, independent of Tks5 expression (Figure 2C). Atelocollagen was also unable to promote invadopodia formation in MDA-MB-231 cells and two other invasive breast cancer cell lines, Hs578t and HCC1806, when compared to native collagen-I (Figure 2C). These data are consistent with a report that fibrillar collagen-I induced an invasive phenotype in breast cancer cells whereas high-density non-fibrillar collagen-I (generated by sonication to shear fibrils) suppressed invasive behavior (Maller et al., 2013).

**Figure 2.**
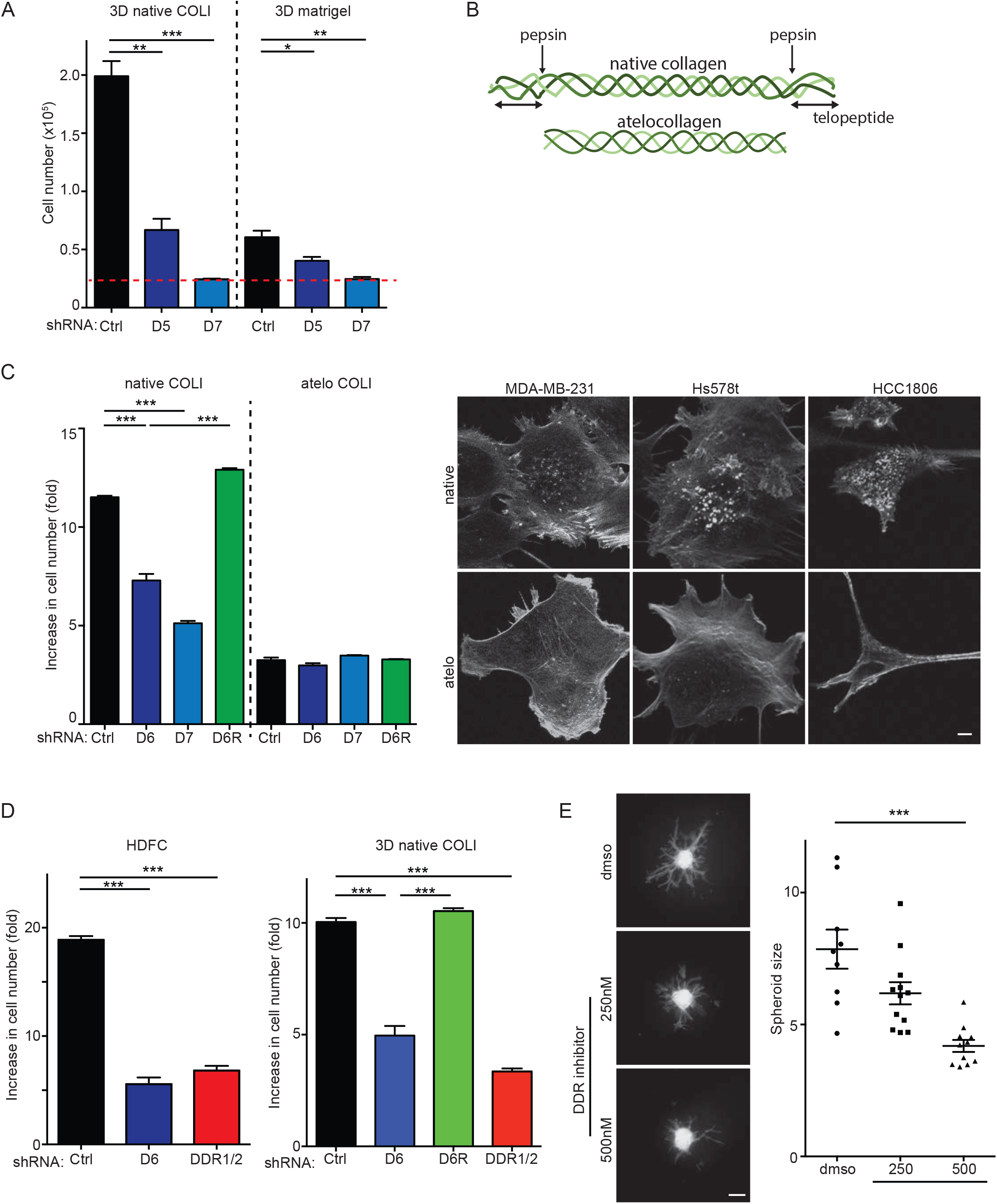
Crosstalk between invadopodia and the extracellular matrix. **A** Growth of cells as indicated in the figure in 3D native type I collagen (3D native COLI) and in 3D matrigel (3D matrigel). Red line shows the cell number when they were embedded in matrix. **B** Schematic view of native collagen and atelocollagen. **C** Growth of cells as indicated in the figure on native type I collagen (native COLI) and atelocollagen (atelo COLI). Fold change in cell number in MDA-MB-231 cells with shRNA-scrambled (Ctrl), shRNA-Tks5 (D6), shRNA-Tks5 (D7) or shRNA-Tks5-D6 + rescued expression of Tks5αGFP (D6R) and representative images (actin by phalloidin) in MDA-MB-231, Hs578t or HCC1806 cells on each conditions (right). **D** Growth of cells as indicated in the figure on high-dense type I collagen (HDFC) and in 3D native type I collagen (3D native COLI). Fold change in cell number in MDA-MB-231 cells with shRNA-scrambled (Ctrl), shRNA-Tks5 (D6), shRNA-Tks5-D6 with rescued expression of Tks5αGFP (D6R) or shRNA-DDR1/2 (DDR1/2). **E** 3D growth/invasion in a hanging droplet spheroid assay with DDR inhibitor, DDR-IN-1. Representative images of spheroids in type I collagen stained by phalloidin (left) and spheroid size measured by actin intensity (right). Scale bars, 5 μm (C) and 500 μm (E). *P*>0.05 unless other specified; *, *P*<0.05; **, *P*<0.01; ***, *P*<0.001.

Epithelial cells interact with collagen via integrins (predominantly α1β1 and α2β1) and the receptor tyrosine kinases DDR1 and DDR2 (Leitinger and Hohenester, 2007), with the DDRs attracting recent attention as possible therapeutic targets (Brakebusch and Fassler, 2005; Rammal et al., 2016; Valiathan et al., 2012). DDR1 is reported to have a kinase-independent function in linearizing invadopodia (Juin et al., 2014). There are no reports on DDR2, although interestingly DDR2 is involved in hypoxia-stimulated invasion of cancer cells (Ren et al., 2014), through stabilization of Snail and subsequent EMT (Zhang et al., 2013). Hypoxia increases invadopodia formation (Diaz et al., 2013), as does Twist-stimulation (Eckert et al., 2011). This suggests a possible reciprocal relationship between invadopodia and collagen-DDR signaling. We used two approaches to investigate this. First, we knocked down DDR1 and DDR2, and tested the effect on growth on and in collagen-I. In both cases we observed a reduction in collagen-I stimulated growth, to the same extent as Tks5 knockdown (Figure 2D). Secondly, we evaluated the effect of the DDR inhibitor DDR1-IN-1, which inhibits both DDR1 and DDR2 at the concentrations used (Kim et al., 2013). We observed a reduction in the invasion and growth of breast cancer cells in 3D (using a spheroid assay), but not 2D growth (Figure 2E, not shown). Together, these data highlight the importance of DDR signaling in invadopodia formation. It is likely that many of these effects occur through DDR2, which is uniquely activated by fibrillar collagen-I (Itoh, 2018; Rammal et al., 2016), whereas DDR1 can be activated by both collagen-I and collagen-IV lacking fibrillar structure. This will be explored in more detail in future work.

### The effect of ECM on invadopodia formation

Research in recent years has highlighted the importance of the tumor microenvironment on invadopodia formation and function, but efforts to systematically evaluate how the microenvironment regulates invadopodia have been hampered by its complexity. Aside from a myriad of insoluble ECM components, the typical tumor microenvironment also consists of soluble growth factors, cytokines and chemokines, secreted by multiple different cell types. With these issues in mind, the Korkola laboratory has developed the microenvironmental microarray (MEMA) technology to enable study of the microenvironment (Watson et al., 2018). MEMA consist of arrayed combinations of ECM molecules printed into each well of a multi-well plate, where each printed ECM spot forms a growth pad upon which cells can be cultured. Each individual well is treated with a separate soluble growth factor or ligand. By performing these assays in a multi-well format, the effects of unique combinations of ECM plus ligand can be assessed for their effects on cellular phenotypes (Watson et al., 2018). We ran a small-scale MEMA experiment using MDA-MB-231 cells in which expression of endogenous Tks5 was suppressed by RNA interference, and replaced by similar levels of Tks5αGFP. We used 45 different ECM proteins, with either no ligand, EGF, FGF or HGF, for a total of 192 different conditions in this screen. After 24 hours of growth, cells were fixed and stained with phalloidin for F-actin visualization and Tks5α co-localization on a confocal microscope (Figure 3). As expected, we observed collagen-I stimulation of invadopodia formation. We also observed stimulation by laminin, consistent with known biology and the observation that invadopodia form at contact sites with basement membrane (Schoumacher et al., 2010). We also saw more invadopodia on fibronectin, an abundant ECM protein with appreciated roles in cancer (Rick et al., 2019). But the most prominent ECM inducer of invadopodia was tropoelastin, the soluble precursor of the cross-linked ECM protein elastin (Vindin et al., 2019). As the name suggests, elastin provides elasticity and resilience to many organs, including the breast and the lung (a frequent site of breast cancer metastasis). Pericellular proteolysis of elastin can impact several aspects of cancer progression (Scandolera et al., 2016), although a discrete role in invadopodia biology has not previously been reported. None of the added growth factors were found to impact invadopodia formation in this MEMA assay format (not shown). This small-scale study demonstrates the feasibility of MEMA to identify novel modulators of invadopodia. In the future, it will be important to also test the roles of their corresponding receptors and downstream signaling events on invadopodia biology.

**Figure 3.**
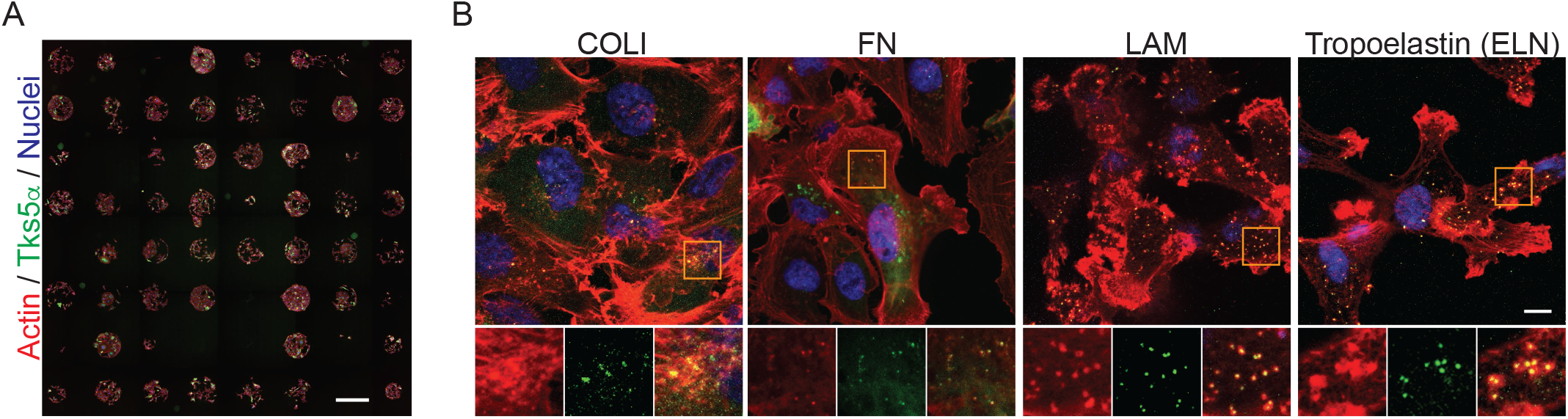
Invadopodia formation assay on microenvironmental microarray (MEMA). **A** Overall view from one of the MEMA assays with MDA-MB-231 cells. Invadopodia were visualized by actin (phalloidin) and Tks5αGFP. **B** Representative zoom-in images from each spot as indicated in the figure (type I collagen: COLI, fibronectin: FN, laminin: LAM and tropoelastin). Magnified areas (orange square at top) shown at bottom. Hoechst staining reveals nuclei of cells in the images. Scale bars, 500 μm (A) and 10 μm (B).

### Spatial distribution of Tks5α and other invadopodia components

Previous studies have started to reveal fundamental properties of invadopodia such as molecular components, order of assembly, and overall architecture. Currently, it is thought that invadopodia formation begins with a ‘core’ of branching cortical actin (with actin, cortactin, N-WASP, cofilin and so on), likely through Src-mediated tyrosine phosphorylation of cortactin, followed by recruitment of SHP2 and Tks5α (Oikawa et al., 2008; Sharma et al., 2013). SHP2 generates PI(3,4)P2, which binds to the PX domain of Tks5α to turn the latter into a multi-functional scaffold to: bind to and stabilize filamentous actin; allow protrusion extension and maturation; engage and activate proteases such as the ADAMs and potentially MMPs for proteolytic activities; and recruit other adaptor proteins such as Nck2 and dynamin-2, which help organize signaling molecules such as the integrins, EGFR, and MET. While recruitment of these proteins and lipids could explain the various biological functions of invadopodia, the prior studies have mostly used 2D culture systems (e.g., on coverslips with or without a thin layer of gelatin). The structure and dynamics of invadopodia in more physiological 3D growth conditions remain poorly defined. We have conducted a series of experiments to define invadopodia in native collagen-I. Unexpectedly, both in tumor cell spheroids in 3D and cells on HDFC, Tks5α localization was restricted at the base of invadopodia but not in the protrusion body itself (Figure 4A, B, E and **Supplementary movie1**). We have also used super-resolution microscopy (SRM) to image Tks5 and the results revealed even more details of Tks5α distribution at the base of invadopodia, resembling that of the actin cortex (Figure 4D and E). Since Tks5 likely acts as a scaffold for multiple proteins, this localization pattern might also confine relevant biological processes to the base of invadopodia.

**Figure 4.**
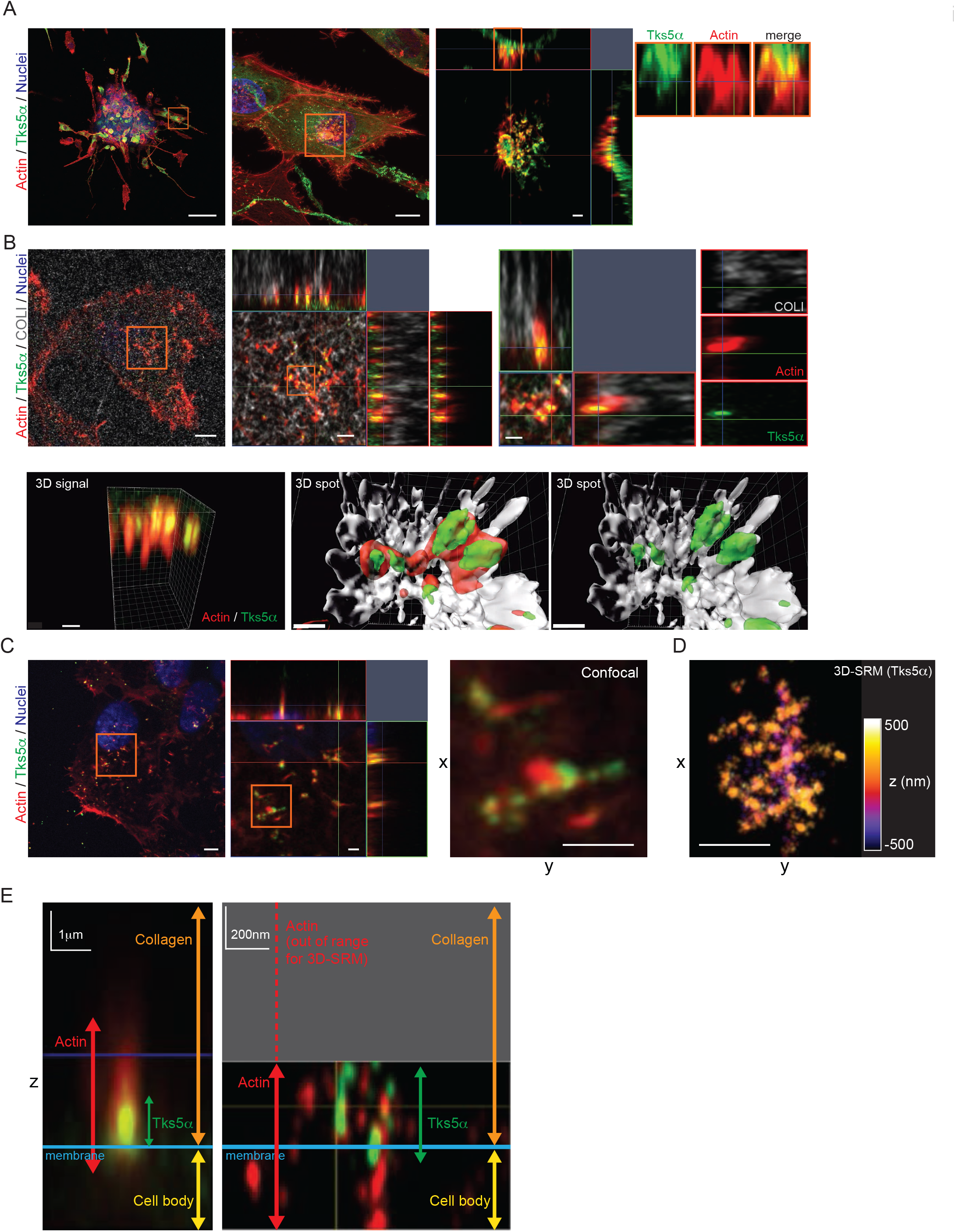
Fine structures of invadopodia in 3D type I collagen. **A** Invadopodia were visualized in a 3D type I collagen hanging droplet spheroids. Representative images of MDA-MB-231 spheroids stained by phalloidin (red) and Tks5αGFP (green). Magnified areas (orange square in left) were shown in middle. Hoechst staining reveals nuclei of cells in the images. Magnified orthogonal view (orange square in middle) shown at right. **B** Invadopodia in MDA-MB-231 cells with Tks5αGFP were visualized on high-dense fibrillar collagen (HDFC). Representative images of invadopodia stained by phalloidin (red), Tks5αGFP (green) and HDFC (gray). Magnified orthogonal view (orange square in left and middle) shown in the middle and atright. 3D reconstruction images were processed by using Imaris software (3D signal or spots, bottom). **C** Invadopodia on HDFC were visualized by phalloidin (red) and Tks5αGFP (green). Hoechst to denote nuclei of cells in the images. Magnified orthogonal view (orange square in left) were shown in middle. Digital zoom-in image from single z-stack plane in middle (orange square) was shown in right. **D** 3D super-resolution microscopy (SRM) image of invadopodia on HDFC. Sample was prepared at same time as in C. **E** Comparison of invadopodia on HDFC between confocal microscopy and SRM. Representative images and structural information as indicated in the figure. Scale bars, 100 μm (A left), 10 μm (A middle), 5 μm (B top left, C left), 2 μm (A right, B top middle) and 1 μm (B top right, B bottom three, C right three).

We next evaluated the spatial distribution of the invadopodia components and actin regulators cortactin, N-WASP and fascin HDFC. To do this, cortactin, N-WASP or fascin-GFP fusion proteins were overexpressed in MDA-MB-231 cells together with Tks5α-mCherry, and the localization of these proteins in invadopodia formed on HDFC evaluated (Figure 5A, B). First, we confirmed that Tks5αGFP and Tks5α-mCherry were colocalized at the base of invadopodia. N-WASP accumulated in similar areas to Tks5α, consistent with reports of their association between N-WASP and Tks5 (Oikawa et al., 2008). However, cortactin, which has also been reported to associate with Tks5α (Crimaldi et al., 2009), extends much further into the invadopodia body than Tks5α. Finally, while there was no localization or accumulation of fascin in invadopodia at day 2 on HDFC, some fascin was detected along the protrusions beginning at day 4. These distributions are schematized in Figure 5C. It will be important to extend these observations to other invadopodia proteins in the future, and to look at the ultrastructure of the F-actin contained in invadopodia over time. However, these observations may suggest that fascin-directed actin cables only form once the protrusions have fully elongated.

**Figure 5.**
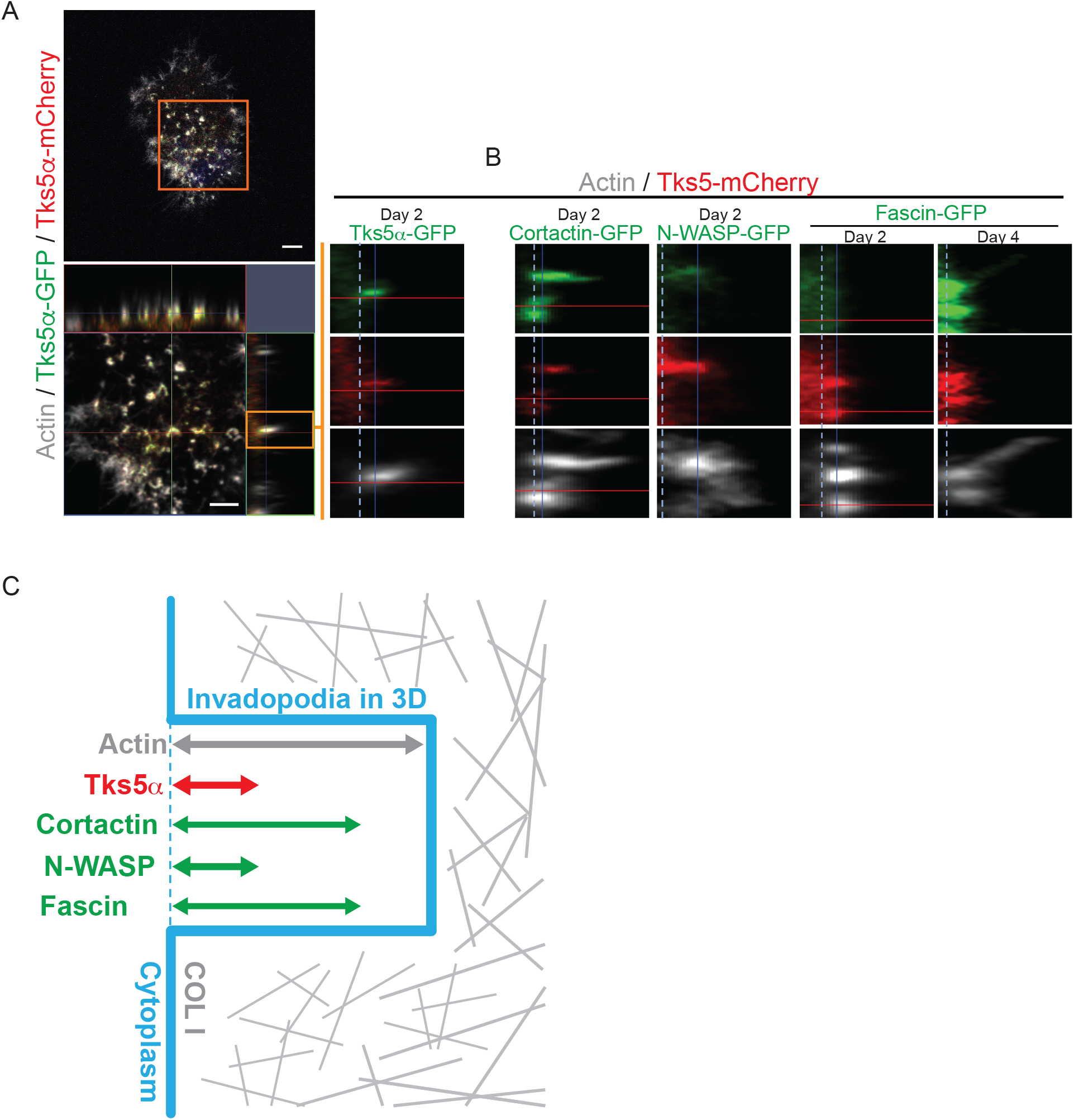
Localization of invadopodia components on HDFC. **A** Invadopodia in MDA-MB-231 cells with Tks5αGFP and Tks5α-mCherry were visualized on HDFC. Representative images of invadopodia stained by phalloidin (gray), Tks5α-mCherry (red) and Tks5αGFP (green). Magnified orthogonal view (orange square in top) shown at bottom. Digital zoom-in side-view image from single invadopodia in bottom (orange square) shown at right. Dot-line shows the area of cell membrane. **B** Invadopodia side-view images in Tks5α-mCherry overexpressing MDA-MB-231 cells with cortactin-GFP, N-WASP-GFP or fascin-GFP on HDFC. **C** Schematic view of invadopodia with localization of invadopodia-related proteins in 3D. Scale bars, 5 μm (A top) and 3 μm (A bottom).

### Localization and function of pericellular proteases

Invadopodia are known to be sites of pericellular proteolysis, with matrix, cysteine and serine protease activity associated with them, although whether protease activity is required for invadopodia formation, or just for function, may be cell type and context dependent (Linder, 2007; Murphy and Courtneidge, 2011). Nor have 3D studies of proteolytic activity associated with invadopodia been performed. We have begun to investigate this by determining the localization of both the MMPs and the cysteine cathepsins, initially focusing on their proteolytically active forms, using activity based probes (ABPs) for cathepsins (Xiao et al., 2013) and MMPs (Saghatelian et al., 2004). In both cases, active enzyme was localized to the base of invadopodia, in approximately the same location as Tks5α in magnified orthogonal views (Figure 6A and C). 3D sideview of invadopodia more clearly showed the spatial localization of those proteins **(Figure6B, 6D, Supplementary movie2 and movie3)**. This was surprising, since it might have been expected that ECM degradation occurs at the advancing tip of the invadopodium. It has been shown that pericellular proteases can be delivered as exosome cargo to invadopodia (Hoshino et al., 2013). In the future, it will be interesting to visualize exosomes and other microvesicles in these 3D systems.

**Figure 6.**
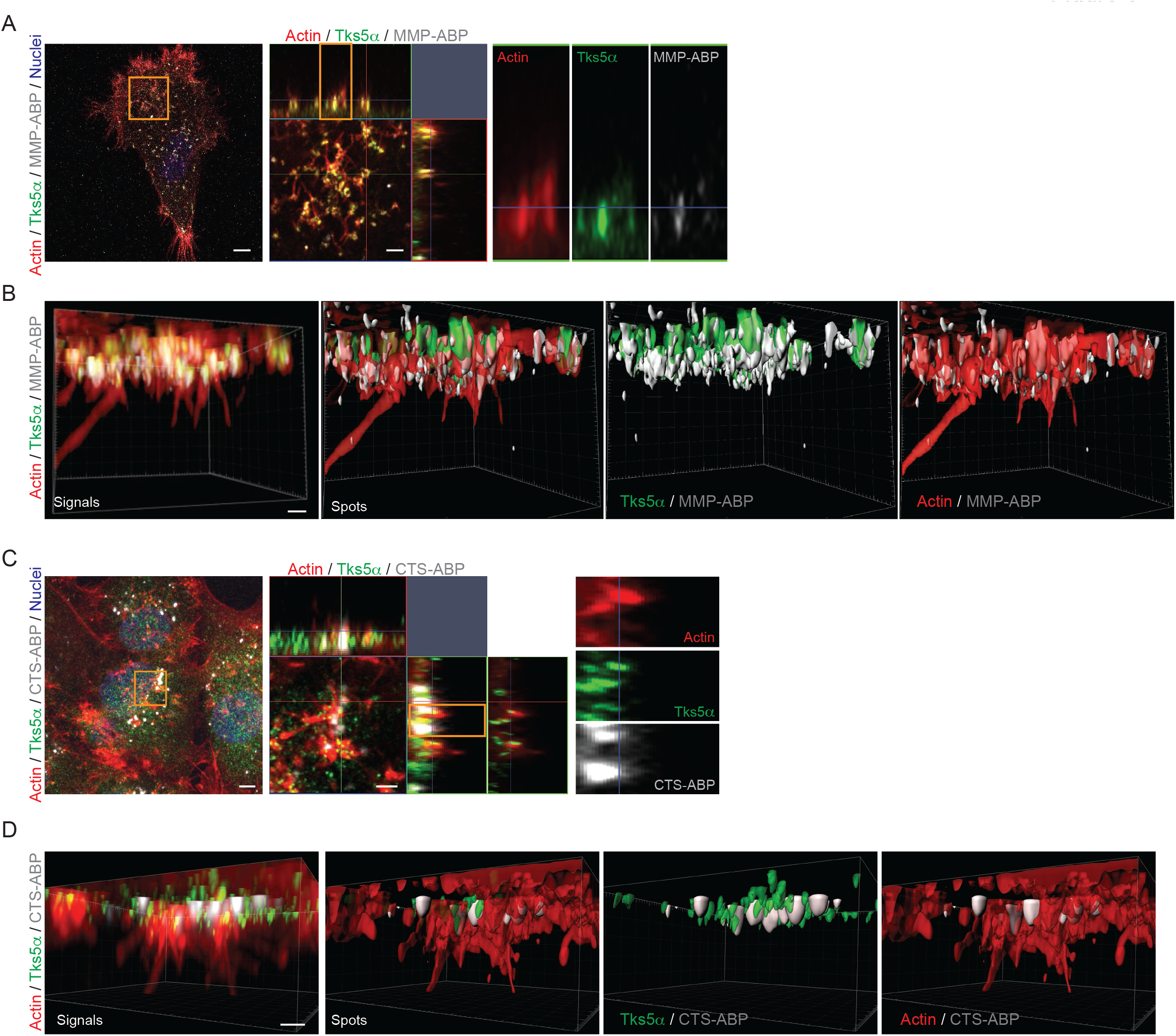
Localization of protease activity at invadopodia on HDFC. **A** MMP activity at Invadopodia in MDA-MB-231 cells with Tks5αGFP were visualized on HDFC. Representative images of invadopodia stained by phalloidin (red), Tks5αGFP (green) and MMP activity (gray, MMP-ABP). Magnified orthogonal view (orange square in left and middle) were shown in middle and right. 3D reconstruction images were processed by using Imaris software (3D signal or spots) and shown in bottom. **B** Cathepsin activity at invadopodia in MDA-MB-231 cells with Tks5αGFP was visualized on HDFC. Representative images of invadopodia stained by phalloidin (red), Tks5α (green, F4 antibody) and cathepsin activity (gray, CTS-ABP). Magnified orthogonal view (orange square in left and middle) shown in middle and at right. 3D reconstruction images were processed by using Imaris software (3D signal or spots) and shown at bottom. Scale bars, 5 μm (A left, C left, D), 2 μm (A right, C right) and 1 μm (B).

We next looked at the requirement for metalloprotease and cysteine cathepsin activity for invadopodia formation, using small molecule inhibitors. We found that 3D growth was affected by the MMP inhibitor GM6001 and the impermeable cathepsin inhibitor CA074 (Figure 7A). These data suggest non-redundant, or concerted, functions for these two classes of protease. We used our high-resolution microscopy techniques to investigate further, comparing to a Src family kinase (SFK) inhibitor SU11333 (Laird et al., 2003), a derivative of SU6656 (Blake et al., 2000), since many invadopodia proteins are Src substrates (Murphy and Courtneidge, 2011). SFK inhibition markedly reduced the Tks5α content at the base of nascent invadopodia, whereas MMP and cathepsin inhibition caused an increase in Tks5α content (Figure 7B and C). We confirmed these observations by plotting signal intensities with z-stack depth (Figure 7C, **right**). SFK inhibition reduced both actin and Tks5α signals at invadopodia (Y-axis), but MMPs/cathepsin inhibition increased Tks5α accumulation at invadopodia compare to the DMSO control group. These graphs also show the decrease of protrusion depth (X-axis, actin intensity) in all inhibitor groups. To test the formation of functional invadopodia on HDFC, the length of invadopodia was measured. We found that all inhibitors markedly reduced the length of protrusions (Figure 7D). Thus, we speculate that SFK signaling may be required for initiation and perhaps maintenance of protrusions, whereas MMPs and cathepsins may promote the maturation/elongation of the structures.

**Figure 7.**
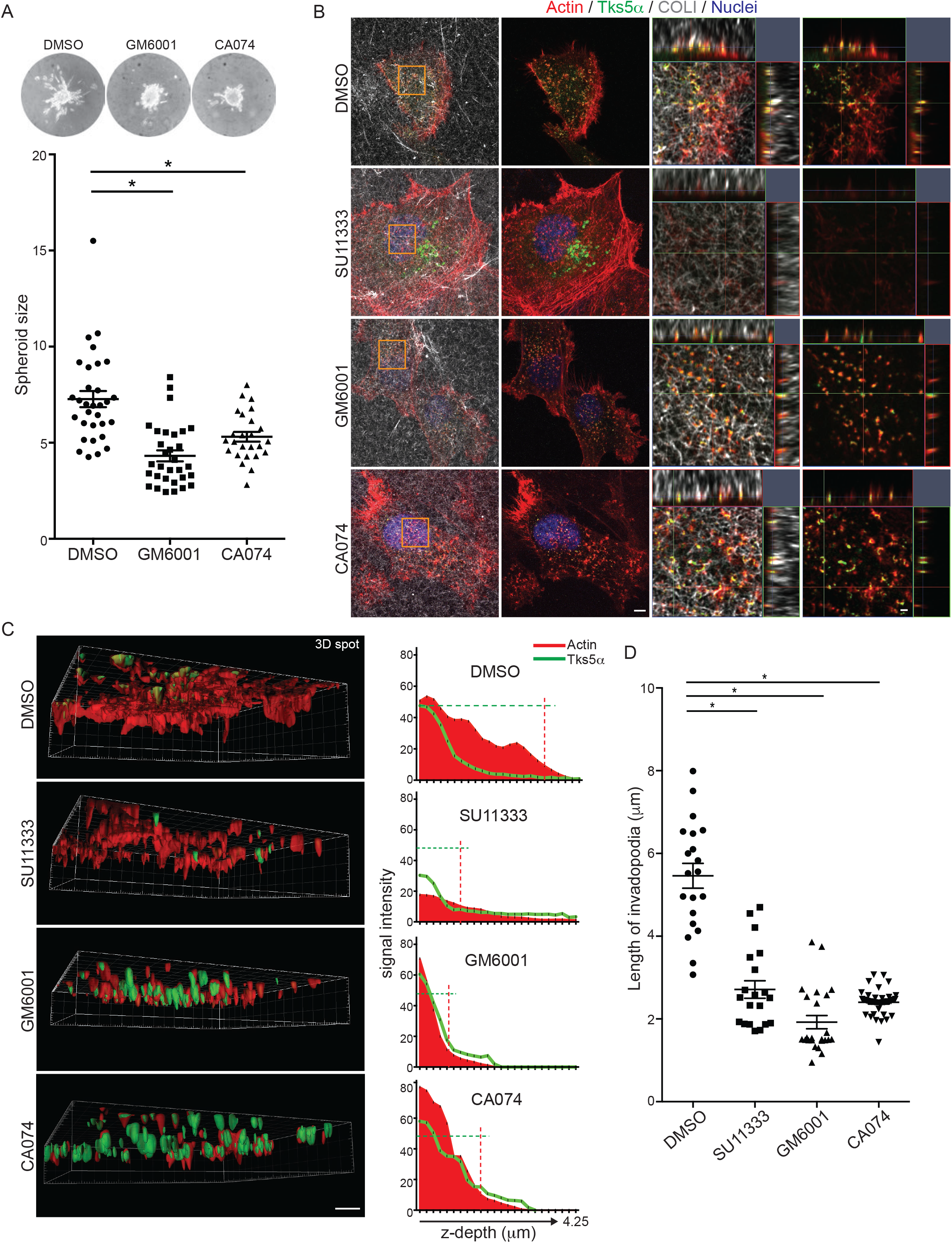
Role of protease activities in 3D invadopodia. **A** 3D growth/invasion in a hanging droplet spheroid assay in MDA-MB-231 cells with inhibitors, GM6001 or CA074. Representative brightfield images of spheroids in type I collagen (top) with spheroid size measured by actin intensity (bottom). **B** Invadopodia in MDA-MB-231 cells with Tks5αGFP were visualized on high-dense fibrillar collagen (HDFC). The cells were treated with DMSO, SU11333 (1μM), GM6001 (10μM) or CA074 (10μM). Representative images of invadopodia stained by phalloidin (red), Tks5αGFP (green) and HDFC (gray). Magnified orthogonal view (orange square) shown in left two panels. **C** Images from B were processed for 3D reconstruction (3D spots) using Imaris software. Length (z-depth) and intensity of actin or Tks5αGFP (from three representative invadopodia) was analyzed in each condition (C, right graphs). Red dotted-line shows depth of actin intensity 10 in each condition. Green dotted-line shows maximum intensity of Tks5αGFP in DMSO. **D** Individual invadopodia length was measured using Imaris software. Scale bars, 5 μm (B left, C) and 1 μm (B right). *P*>0.05 unless other specified; *, *P*<0.001.

## CONCLUSIONS

In conclusion, we describe here two new antibodies: one (G6) recognizes all isoforms of Tks5; and one (F4) is specific for Tks5α. Both have antigen recognition utility in multiple formats including immunoblotting, immunoprecipitation and immunostaining (both immunohistochemistry and immunocytochemistry), making them highly valuable to the invadosome community. Combining these antibodies with several cutting-edge technologies such as super-resolution microscopy, MEMA and activity-based probes, we have begun to investigate the structure and function of invadopodia forming in response to the ECM. The results revealed new and complex sub-invadopodia architecture unseen in our previous analyses in 2D, and ECM components (tropoelastin) as novel invadopodia inducer. Among others, it is possible to evaluate whether invadopodia architecture is influenced by the ECM components present by studying invadopodia structure in 3D or semi-3D model systems. The 3D and HDFC formats also revealed that ECM stimulates cancer cell growth in an invadopodia-dependent manner, which goes a long way to explaining why Tks5α (and other invadopodia proteins) are required for growth of many invasive cancer cells in 3D, but not when the same cells are cultured on tissue culture plastic. Future studies will expand on the findings described herein, with the goals being to evaluate more invadopodia components, to map their localizations, interactions and functionality. These studies will be important to determine whether invadopodia initiators such as the DDR receptors co-localize to invadopodia, as well as the spatial localization of their downstream signaling components.

## ACKNOWLEDGMENTS

We acknowledge the expert assistance of Dr. Stefanie Kaech Petrie and Crystal Chaw (cell imaging) in the Advanced Multiscale Microscopy Shared Resource, Dr. Philip R Streeter and YongPing Zhong (G6 antibody production) Oregon Stem Cell Center, Monoclonal Antibody Core, and Dr. David Kilburn (MEMA preparation) in the Department of Biomedical Engineering, Knight Cancer Institute at the OHSU. We also thank Dr. Ting Zheng for assistance with conjugating antibodies used in DNA-PAINT experiments.

This work was supported in part by the National Institutes of Health grant R01 CA217625 and support from the Knight Cancer Institute (SAC), Brenden Colson Center for Pancreatic Care at OHSU (SLG).

