## Supplementary figure for "Crosstalk between invadopodia and the extracellular matrix"

A

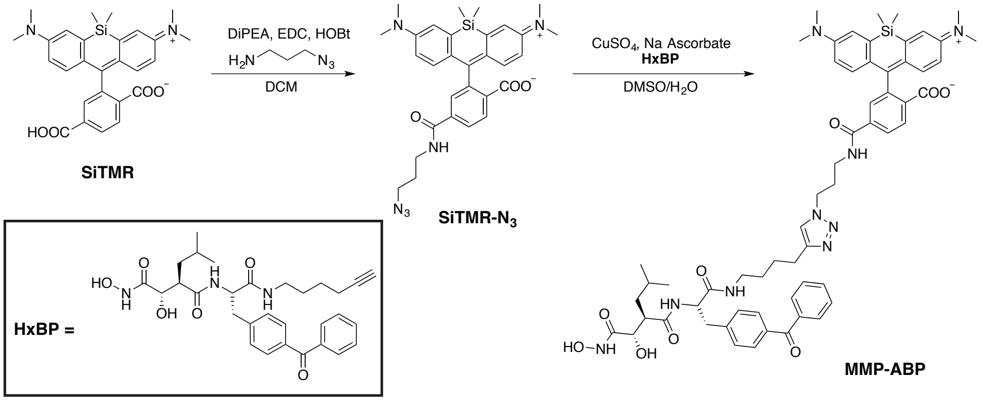


B

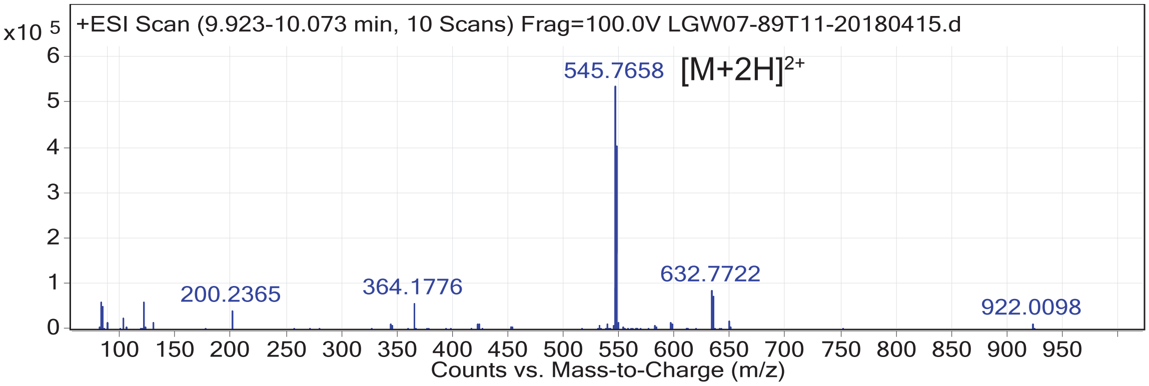


C

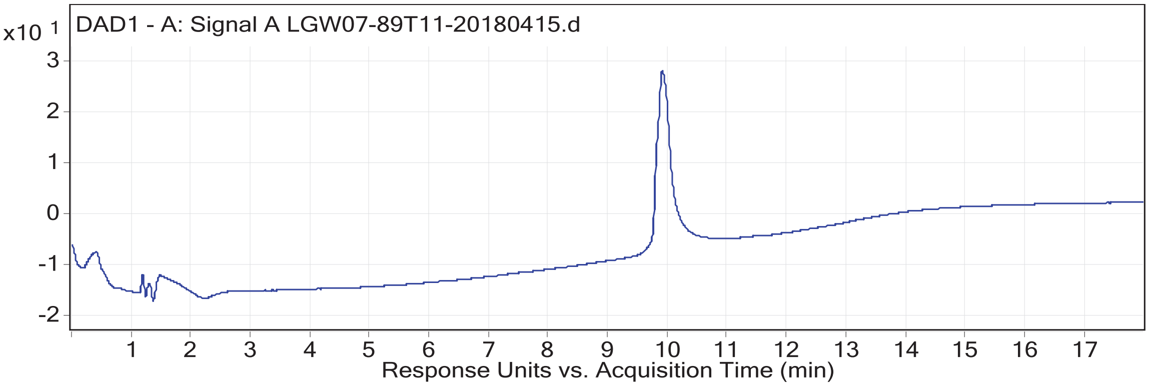


**Supplementary Figure S1. Chemical synthesis of matrix metalloproteinase activity-based probe (MMP-ABP). A**. Synthetic route to the MMP-ABP labeled with SiTMR using click chemistry. DiPEA = N,N-diisopropylethylamine; EDC = N-(3-Dimethylaminopropyl)-N′-ethylcarbodiimide hydrochloride; HOBt = 1-Hydroxybenzotriazole hydrate; DCM = dichloromethane; DMSO = dimethyl sulfoxide. **B** structure and **C** purity confirmation was obtained using HPLC-MS via analysis of mass-to-charge (m/z) ratio in positive ion mode and absorbance at 254 nm.
